# Development and validation of a mouse model of contemporary cannabis smoke exposure

**DOI:** 10.1101/2021.02.11.430865

**Authors:** Matthew F. Fantauzzi, Steven P. Cass, Joshua J.C. McGrath, Danya Thayaparan, Peiyao Wang, Martin R. Stampfli, Jeremy A. Hirota

**Author notes:** **Corresponding Author:** Jeremy Hirota, PhD, Firestone Institute for Respiratory Health – Division of Respirology, Department of Medicine, McMaster University, Hamilton, ON, L8N 4A6.

## Abstract

Cannabis is widely used for both recreational and medicinal purposes. Inhalation of combusted cannabis smoke is the most common mode of drug consumption, exposing the lungs to the pharmacologically active ingredients, including tetrahydrocannabinol (THC) and cannabidiol (CBD). While the relationship between cannabis smoke exposure and compromised respiratory health has yet to be sufficiently defined, previous investigations suggest that cannabis smoke may dysregulate pulmonary immunity. Presently, there exists few pre-clinical animal models that have been extensively validated for contemporary cannabis smoke exposure. To address this need, we developed a mouse model with readouts of total particular matter, serum cannabinoid and carboxyhemoglobin levels, lung cellular responses, and immune mediator production.

Using a commercially available smoke exposure system and a cannabis source material of documented THC/CBD composition, we exposed mice to a total particulate matter of 698.89 (SD = 66.09) µg/L and demonstrate increases in serum cannabinoids and carboxyhemoglobin. We demonstrate that cannabis smoke modulates immune cell populations and mediators in both male and female BALB/c mice. This modulation is highlighted by increases in airway and lung tissue macrophage populations, including tissue-resident alveolar macrophages, monocyte-derived alveolar macrophages, and interstitial macrophage subpopulations. No changes in airway or lung tissue infiltration of neutrophils were observed. Immune mediator analysis indicated significant upregulation of MDC, TARC, and VEGF within the lung tissue of cannabis smoke exposed mice. This accessible and reproducible smoke exposure model provides a foundation to explore the impact of chronic cannabis exposures and/or co-exposures with pathogens of clinical relevance, such as influenza.

## INTRODUCTION

Cannabis is a mood-altering drug that is widely used for its psychoactive and remedial effects. It is the most commonly consumed drug worldwide with an estimated 192 million users – a number which is expected to increase due to trends in recreational legalization as well as innovations in medical applications^1^. The drug is comprised of over 400 biologically active compounds with ∼25% of these being cannabinoids, most notably tetrahydrocannabinol (THC) and cannabidiol (CBD)^2^. These compounds are thought to play roles in a variety of physiological processes including neurocognition, appetite stimulation, pain alleviation, and immunity^3–6^. Notably, the method by which cannabis is consumed can both influence the pharmacological response to the drug and modulate the risk of collateral health effects. Among these routes, inhalation of combusted cannabis smoke is the most common mode of drug consumption, exposing the lungs to the pharmacologically active ingredients as well as a plethora of combustion products^7,8^. Research investigating the potential health risks of inhaled smoke is highly warranted given our insufficient understanding of the biological consequences of cannabis smoke exposure.

Much like tobacco cigarette smoking, cannabis smoke exposure has been implicated in the development of adverse respiratory outcomes. Cannabis smoking is linked to respiratory symptomology including an increased incidence of cough, wheezing, chest tightness, and shortness of breath^9–12^. Studies also demonstrate a potential link between cannabis use and an increased risk of pulmonary infection, suggesting that cannabis smoke may compromise respiratory host defense^13–16^. Although there is currently no strong association between cannabis smoking alone and the development of chronic obstructive pulmonary disease (COPD), research indicates a potential synergistic effect between cannabis and tobacco smoke that may exacerbate lung inflammation and disease development^17,18^. While work delineating the exact mechanisms by which cannabis smoke promotes lung disease pathogenesis is limited, several studies have implicated dysregulated pulmonary immunity as a factor in these processes^19–21^. Despite this, limited experimental models are available to perform detailed assessments of how cannabis smoke exposure directly contributes to modulation of lung immunity.

While rodent models have been used in-depth to characterize tobacco smoke-associated lung inflammation, cannabis smoke studies using similar approaches remain largely unexplored. Historical *in vivo* investigations utilizing cannabis smoke exposure systems require an update as the composition of cannabis has varied significantly over time and since legalization^22,23^. The average concentration of THC in a commercially available strain can exceed 15%, whereas earlier models used strains with THC levels below 5%. Consequently, early research on cannabis smoke may not accurately reflect the health implications of modern cannabis use. To date, few rodent smoke exposure studies using compositionally relevant strains of cannabis exist^24^, particularly those procured from legal suppliers. This omission, in conjunction with evolution in the commercial cannabis landscape, support the need for further investigations characterizing the mechanisms through which cannabis smoke contributes to lung inflammation and pathology.

When considering how cannabis smoke exposure modulates immunological processes in the lungs, once can consider the well-characterized effects of tobacco cigarette smoke on pulmonary inflammation. Supporting this comparison are the chemical compositional similarities between the smoke of each combusted plant product, which suggests that any inflammatory effects may be conserved^8^. Studies investigating tobacco smoke exposure have identified key changes to immune cell population dynamics and function. Altered innate immunity has been suggested to be an early marker and inducer of smoke-associated disruption to lung homeostasis. In particular, it has been demonstrated that lung macrophages and neutrophils perform an essential role in mediating tobacco smoke-induced inflammation^25^. The central paradigm suggests that proinflammatory changes to lung macrophage function and quantity leads to pulmonary neutrophilia and subsequent exacerbated inflammation following tobacco smoke exposure^26,27^. Previously, this paradigm centered around the role of tissue-resident alveolar macrophages, however, unexplored until recently macrophage subtypes such as monocyte-derived alveolar macrophages and interstitial macrophage populations 1 – 3 (IM1, IM2. IM3) have been hypothesized to play a role as well^28,29^. While these populations have yet to be shown to be involved in tobacco smoke-associated immunomodulation, their suggested contribution to proinflammatory outcomes in the lung supports their inclusion as possible effector cells in smoke-induced inflammation^29,30^. The impact of cannabis smoke exposure on these macrophage populations and other immune cells potentially important in smoke-associated lung inflammation remains to be extensively unexplored.

In the present study, we develop a mouse model of contemporary cannabis smoke exposure using a legal source of defined composition. We confirm that our exposure system delivers cannabis smoke to mice using readouts of serum cannabinoids and carboxyhemoglobin. Using a four day smoke exposure model, we saw the induction of changes to lung immune cellularity and mediator levels, suggesting cannabis smoke is not innocuous. Importantly, we establish and validate the system using commercially available cannabis strains, void of potential confounding contaminants, that is consistent with relevant compositions available legally worldwide.

## METHODS

### Animals

Six to eight week old male and female BALB/c mice were purchased from Charles River Laboratories (Quebec, Canada). Mice were housed in the McMaster Central Animal Facility with *ad libitum* access to food and water and subjected to a 12-hour lighting cycle. All experimental procedures were approved by the Animal Research Ethics Board of McMaster University.

### Cannabis cigarette preparation

Cannabis cigarettes were prepared using cannabis purchased from the Ontario Cannabis Store (Ontario, Canada). All strains used were indica dominant and contained 10 – 14% THC and 0 – 2% CBD. Cigarettes were hand-rolled by grinding dried cannabis flower, packing into king size Premier cigarette tubes (R.J. Reynolds Tobacco Company, USA), and twisting off the end to seal the cigarette (Fig. 1A, 1B). Each cigarette contained 0.84 (± 0.06) grams of cannabis. Prior to running the smoke exposure protocol, the filter of each cigarette was removed, and the cannabis was packed towards the closed end (Fig. 1C).

**Figure 1.**
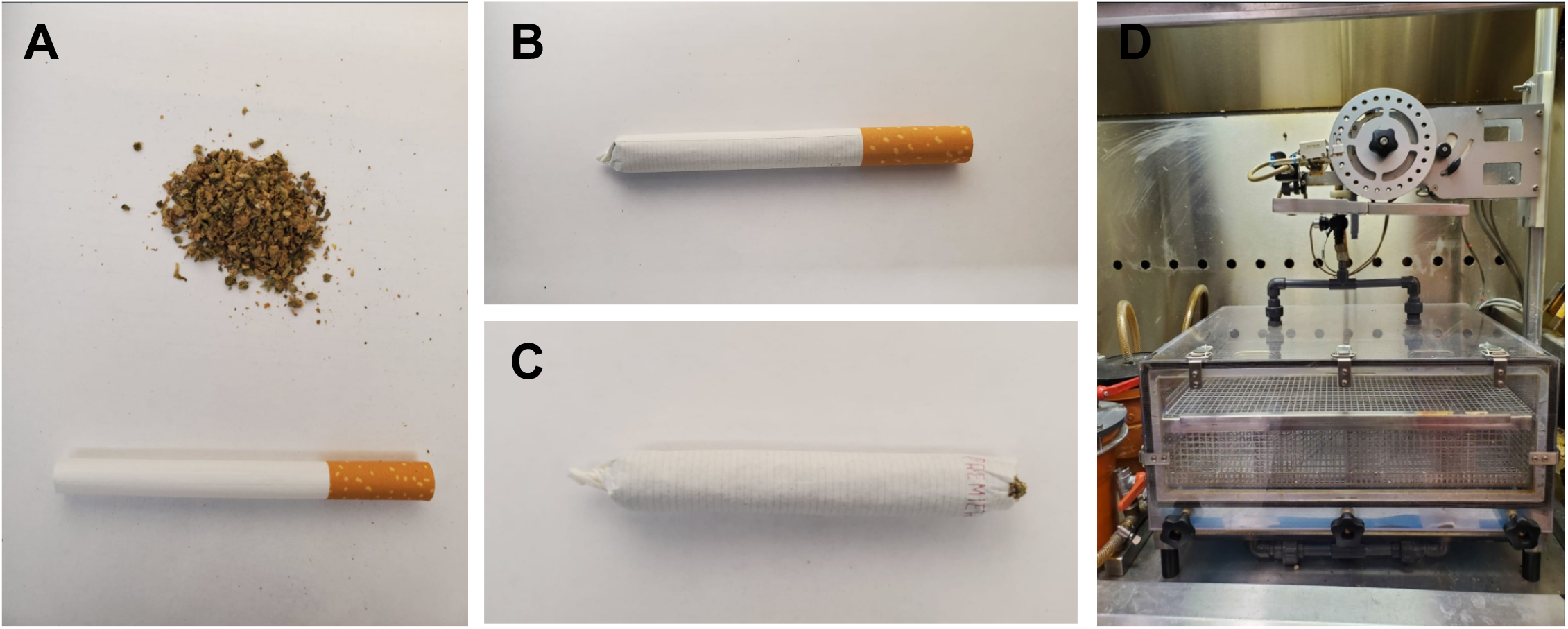
Cannabis cigarette preparation and smoke exposure system. Cannabis cigarettes are prepared by (A) grinding 0.84 (± 0.06) grams of dried cannabis flower (10 – 14% THC, 0% CBD), (B) packing into king size empty cigarette tubes and twisting off the end to seal the cigarette. (C) The filter of each cigarette is removed, and cannabis is packed towards the sealed end of the cigarette prior to smoke exposure. (D) Cannabis cigarettes are loaded into the cigarette holder on a Smoke Inhalation Unit 24 system. Experimental animals are placed in the lower chamber. Cannabis smoke and fresh air are pumped into the chamber following a computerized protocol attached to the system.

### Cannabis smoke exposure protocol

Prior to cannabis smoke exposure, mice were acclimatized to restrainers over three days. Mice were whole-body exposed using a Smoke Inhalation Unit 24 (Promech, Sweden) (Fig. 1D) to the smoke of six cannabis cigarettes twice daily for one day or four days, depending on the experimental design. Mice were given a three hour break in between daily exposure sessions. Control mice were sham-exposed to room air.

### Total particulate matter (TPM) measurement

TPM was measured in the smoke exposure system using a Flowmeter 4045 (TSI Incorporated, USA) and paper filters. Six cigarettes were placed in the exposure system and burned using the same settings as the experimental smoke exposure procedure. Smoke and air outflow was measured during the burning of the first cigarette using the Flowmeter 4045. Filters were placed and subsequently removed one at a time in the outflow tubing during the burning of cigarettes one, three, and five. Following the exposure, the filters were weighed and TPM was derived by dividing the average change in filter mass by the average change in outflow multiplied by four.

### Carboxyhemoglobin and cannabinoid measurement

Fifteen minutes or one hour following a smoke exposure period, whole blood was collected from smoke exposed and control mice using capillary tubes via retro-orbital bleed. Heparinized capillary tubes were used to collect whole blood from the opposite eye. Carboxyhemoglobin in whole blood samples was quantified via CO-oximetry by the McMaster University Medical Centre core laboratory. Plasma was collected by incubating and centrifuging heparinized whole blood samples. Samples were prepared for THC, carboxy-THC, and CBD quantification via mass spectroscopy by the McMaster University Centre for Microbial Chemical Biology. Briefly, samples were treated with cold acetonitrile containing 1% (v/v) formic and 50 ng/mL of internal standard mixture. Next, HPLC grade water was added to the samples, which were then injected into an Agilent 1290 Infinity II HPLC coupled to an Agilent 6495C iFunnel QQQ mass spectrometer (Agilent, California, USA). Separated analytes were eluted and introduced to the mass spectrometer. Quantitation of each cannabinoid was based on the peak area measurement measured by the mass spectrometer. Pooled blank mouse plasma to be used as matrix background was processed using the same protocol as the samples. Standards of THC, carboxy-THC, and CBD were purchased (Sigma Aldrich Canada, Ontario, Canada), prepared in the matrix background, and processed using the same protocol as the samples.

### Bronchoalveolar lavage collection and cell differential quantification

Following four days of cannabis smoke exposure, mice were anesthetized with isoflurane and euthanized via asphyxiation and cardiac puncture. Whole lungs and trachea were excised from the chest cavity. Bronchoalveolar lavage (BAL) was collected by cannulating the trachea, tying off the right lung lobe, instilling the left lobe with 250 µL of cold sterile phosphate buffered saline (PBS), and re-aspirating the fluid. Two sequential instillations were performed in this manner and combined. BAL samples were centrifuged, supernatant removed, and resuspended in 130 µL of PBS. Cell counts were performed on the resuspended samples via haemocytometer and light microscopy. Remaining samples were used for cell differentials via cytospins. BAL samples were centrifuged onto pre-wet microscope slides and then stained using Hema-3 fixative and solutions. Mononuclear and polymorphonuclear cell populations were identified and imaged using Zen microscopy imaging software (Zeiss International, Germany). Cell differentials were multiplied by cell counts to derive differential cell counts.

### Lung tissue processing and flow cytometric analysis

Following instillation of the left lobe for BAL collection, the right lung lobe was excised and collected in RPMI. Collected lobes were incubated for one hour at 37°C in 10 mL of cRPMI with 1500 units of 260 units/mg type 1 collagenase (Worthington Biochemical, Cat. #LS004194) per lobe and pressed through a 35 µm filter. Samples were centrifuged and treated with Ammonium-Chloride-Potassium (ACK) lysis buffer. Following centrifugation, samples were resuspended in 1% bovine serum albumin with 2mM EDTA. Aliquots of the single cell suspension were used for cell counts as outlined in the previous methods section. Single cell suspensions were plated, centrifuged, treated with blocking solution, and stained with fluorophore-labelled antibodies. Samples were then assessed for immune cell population composition and quantity using a BD LSRFortessa Flow Cytometry (BD Biosciences, Canada). The following antibodies were used for flow cytometric analysis (Supp. Fig. 1): LIVE/DEAD™ Fixable Yellow Dead Cell Stain Kit (ThermoFisher Scientific, Cat. #L34967), CD45-Alexa Fluor 700 (BioLegend, Cat. #103127), B220-APC-Cy7 (BioLegend, Cat. #103211), CD3e-APC-Cy7 (BioLegend, Cat. #100329), EpCAM-APC-Cy7 (BioLegend, Cat. #118217), Ly6G-BV785 (BioLegend, Cat. #127645), CD11b-PE-Dazzle 594 (BioLegend, Cat. #101255), CD64-PE-Cy7 (BioLegend, Cat. #139313), MerTK-APC (BioLegend, Cat. #151507), SiglecF-PE (BioLegend, Cat. #155505), CD11c-BV650 (BioLegend, Cat. #117339), CD24-BV421 (BioLegend, Cat. #101825), MHCII-PerCP-Cy5.5 (BioLegend, Cat. #107625), and Ly6C-BV510 (BioLegend, Cat. #128033).

### Immune mediator quantification

Left lung lobes were collected following BAL instillation. Lobes were homogenized in PBS, centrifuged, and supernatant was collected. Immune mediators were quantified in lung tissue homogenate supernatant using the Mouse Cytokine Array / Chemokine Array 44-plex (Eve Technologies, Calgary, Alberta, Canada). Point to point semi-logarithmic analysis was applied to all immune meditator quantities.

### Statistical analysis

GraphPad Prism 9 (GraphPad Software Inc., USA) was used for statistical analyses. The data was expressed in terms of mean and standard errors of the mean (SEM). Unpaired t-tests were performed to compare the means of two groups within the same sex. Differences were considered statistically significant when p < 0.05.

## RESULTS

### Cannabis smoke combustion, inhalation, and systemic distribution

We first sought to establish a mouse model of cannabis smoke exposure using a Smoke Inhalation Unit 24 which has previously been used in tobacco smoke exposure studies (Fig. 1). Six cannabis cigarettes were selected for each exposure session based on preliminary experiments that qualitatively assessed animal behaviour and tolerance (data not shown). Combustion was quantified by total particulate matter (TPM) and carried out in a single smoke exposure session. During exposure, the average TPM concentration within the exposure chamber was 698.89 (SD = 66.09) µg/L. Cannabis smoke exposed mice had reduced levels of activity and ventilation rates (data not shown). Upon removal from the chamber, mice appeared lethargic and hunched but returned to normal activity levels and posture within one hour. Despite these behavioural changes, cannabis smoke exposure was well tolerated with no reduction in clinical score (data not shown). For comparative purposes, these changes in behaviour are also observed in our well-characterized tobacco cigarette smoke models using the same exposure system^25,26^.

Following removal from the smoke exposure chamber, blood was collected within 15 minutes for quantification of metabolites of combustion and phytocannabinoids. Carboxyhemoglobin (COHb), a metabolite produced when hemoglobin is bound by carbon monoxide inhaled from combustion, was significantly elevated in both male and female smoke exposed mice (Fig. 2A). The elevation in COHb returned to baseline levels 60 minutes post-exposure in both males and females (Supp. Fig. 2A). To validate phytocannabinoid delivery through smoke exposure, plasma samples were analyzed via mass spectroscopy (Fig. 1B). Plasma THC and carboxy-THC were both present in all male and female smoke exposed mice compared to controls, which had no traces of either cannabinoid detected. CBD was not detected in any tested samples, consistent with the composition of the selected cannabis strains (reported as 10 – 14% THC and 0 – 2% CBD). In plasma samples collected 60 minutes post exposure, THC was detected in 1/5 males and 1/5 females, while carboxy-THC was detected in 3/5 males and 3/5 females (Supp. Fig. 2B). Plasma samples with detectable THC and carboxy-THC had levels only slightly above the lower level of detection for the analysis. Similar to the earlier assessment, both CBD was undetected in all samples at 60 minutes post exposure.

**Figure 2.**
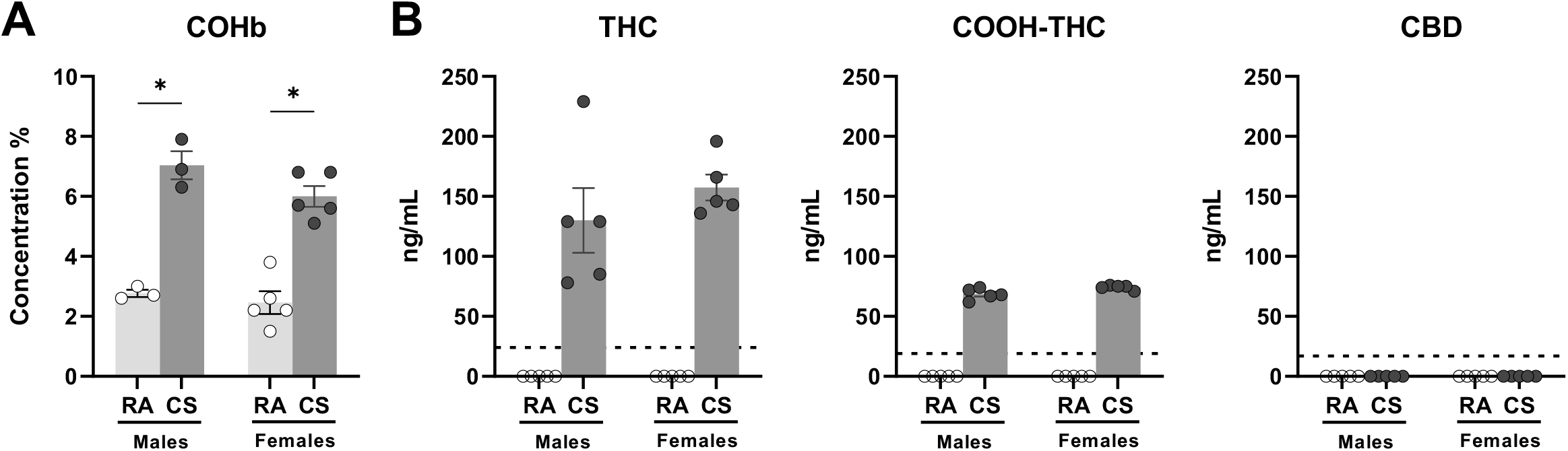
Acute cannabis smoke exposure increases plasma carboxyhemoglobin, THC, and carboxy-THC. Male and female 6 – 8 week old BALB/c mice were exposed to room air (RA) or the smoke of 6 cannabis cigarettes (CS) twice in a day. Whole blood was collected via retro-orbital bleed within 15 minutes following the second exposure session. (A) Carboxyhemoglobin (COHb) percentage was quantified via CO-oximetry. (B) Plasma tetrahydrocannabinol (THC), cannabidiol (CBD), and carboxy-tetrahydrocannabinol (COOH-THC) were quantified via mass spectroscopy. Dotted lines represent the limit of detection for specific cannabinoid. Data points at 0 ng/mL represent values below the limit of detection in cannabinoid analysis. Data represent mean ± SEM; n = 3 – 5/group; *P<0.05, unpaired t-test within each sex in COHb analysis.

Collectively, our results demonstrated that a commercially available smoke exposure system is able to combust cannabis material filled into pre-formed cigarette tubes, resulting in qualitative behaviour changes and systemic cannabinoid distribution consistent with expected metabolite kinetics.

### Cannabis smoke exposure modulates immune cell populations in the airways

Next, we characterized the effects of cannabis smoke exposure on respiratory immune cell populations and immune mediators using a four day model. Total immune cellularity was significantly increased in female cannabis smoke exposed BAL samples compared to control (Fig. 3) Differential cell analysis demonstrated that this increase was driven by an expansion of mononuclear cells; no polymorphonuclear cells were present in any of the female samples. Total immune cellularity and mononuclear cell quantities did not significantly change in male cannabis smoke exposed BAL samples.

**Figure 3.**
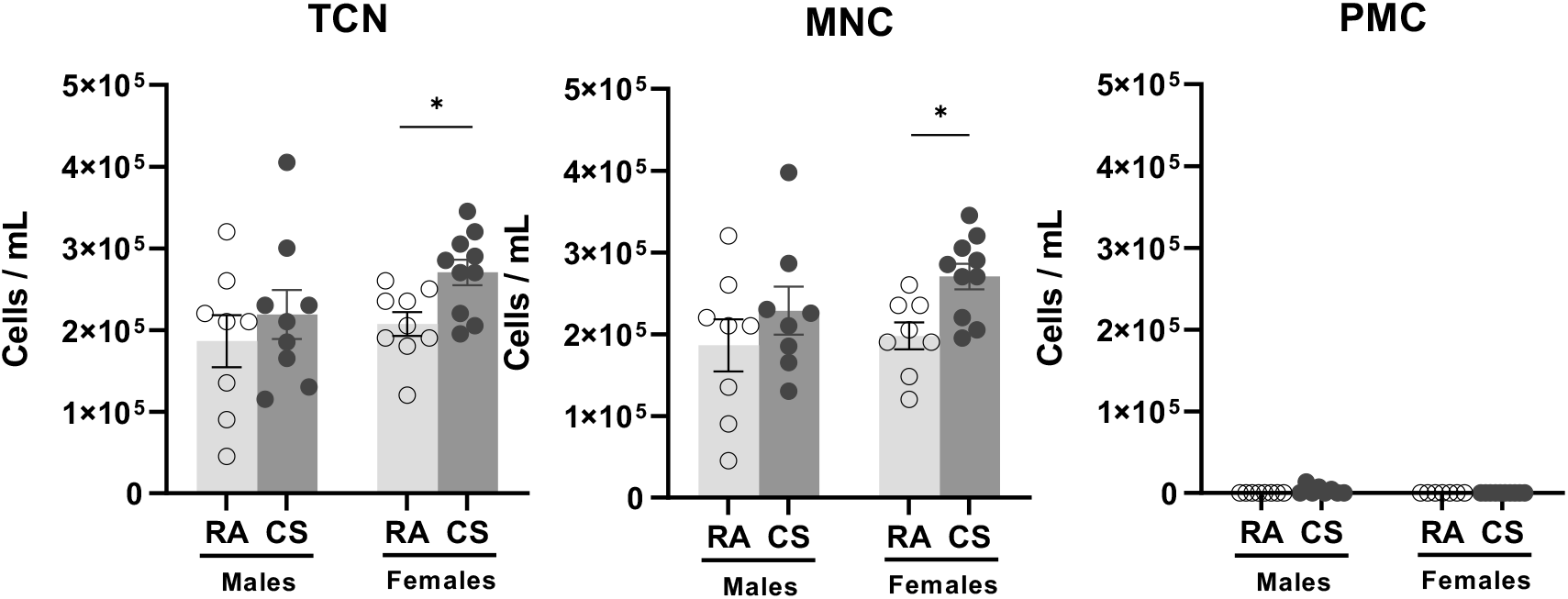
Cannabis smoke exposure modulates cellularity in the bronchoalveolar lavage. Male and female 6 – 8 week old BALB/c mice were exposed to room air (RA) or the smoke of 6 cannabis cigarettes (CS) twice a day for 4 days. (A) BAL total cell number (TCN), mononuclear cells (MNC), and polymorphonuclear cells (PMC) were determined via haemocytometer and cytospin differentials. Data represent mean ± SEM; n = 8 – 10/group, data pooled from two identical experiments; *P<0.05, unpaired t-test within each sex.

### Cannabis smoke exposure modulates immune cell populations in the lungs

Flow cytometric analysis of lung tissue samples further demonstrated modulation of immune cell populations with some distinct differences between sexes. The proportion of total CD45^+^ in the lungs remained unchanged in cannabis-exposed animals, regardless of sex. However, in female cannabis-exposed mice, the proportion of eosinophils were significantly increased; total macrophages, neutrophils, and monocytes were unchanged; and dendritic cells were significantly decreased. In male cannabis-exposed mice, the proportion of macrophages were significantly increased; neutrophils, dendritic cells, and eosinophils were unchanged; and monocytes were significantly decreased.

Specific lung macrophage subpopulations have been suggested to be important in the development of lung inflammation^28,30^. We therefore included complementary cell markers in our flow cytometric analysis to identify relevant subtypes (Fig. 5). In female cannabis-exposed mice, only the proportion of IM1s were significantly increased; all other subpopulations were unchanged. In male cannabis-exposed mice, tissue-resident alveolar macrophages, monocyte-derived alveolar macrophages, and IM1s were significantly increased while IM2s and IM3s were unchanged as a proportion of total leukocytes, compared to control. While most subpopulations were detected in all samples, population size as a proportion of total leukocytes was near-zero for some samples, notably within the monocyte-derived alveolar macrophage and IM3 subpopulations.

Overall, our results demonstrate that a four day cannabis smoke exposure protocol results in modest changes in immune cell populations within the airways and lungs. Notably, while trends are largely conserved between female and males, lung macrophage populations are differentially affected by cannabis smoke exposure.

### Impact of cannabis smoke exposure on immune mediators in the lungs

To perform a broad survey of the impact of cannabis smoke exposure on lung immune responses, we performed a 44-plex cytokine array on homogenized lung tissue (for all data see Supp. Table 1). IL-1α, an interleukin-1 family cytokine which has been demonstrated to be involved in smoke-associated inflammation^27^, was unchanged in cannabis smoke exposed mice compared to control (Fig. 6A). Detectable macrophage-associated immune mediators such as MCP-5, MIP-2, and MIP-3β were also unchanged, while macrophage-derived chemokine (MDC) was significantly increased in male smoke exposed mice (Fig. 6B). Anti-viral cytokines, which are typically involved in early innate immune signaling in response to viral infection and have been proposed to be suppressed by cannabis smoke exposure^31,32^, such as IFNβ-1, IP-10 (CXCL10), and RANTES (CCL5), were unchanged in cannabis smoke exposed mice compared to control (Fig. 6C). Additional cytokines including CCL17 (TARC) in the males (p = 0.037) and VEGF in the females were significantly increased compared to room air control (Fig. 6D), with trends conserved across sexes.

In summary, immune profiling in lung tissue following cannabis smoke exposure reveals elevations in TARC and VEGF, with limited impact on IL-1 family cytokines, macrophage and monocyte chemotactic factors, and antiviral mediators.

## DISCUSSION

The increasing popularity of smoking cannabis recreationally and medicinally has brought to light the absence of experimental evidence of the potential lung health risks associated with inhalation. Given that our limited knowledge is based on historical data where cannabis compositions significantly varied, we sought to understand the affect of inhaled cannabis smoke using a well-controlled system and compositionally relevant cannabis strains^22,23^. We demonstrated that cannabis smoke exposure resulted in systemic distribution of cannabinoids and associated behaviour changes. Lung immune profiling reveals consistent patterns between male and female mice, with changes in monocyte and macrophage cell populations and immune mediators. Our results form a modern foundation for further exploration of different strains of cannabis, exposure protocols, and interaction with pathogens important in lung health and disease.

The development of this cannabis smoke exposure model was based heavily on existing models of tobacco smoke exposure that have been successfully used in respiratory investigations, among others^25^. To note, cannabis provides unique challenges in terms of model development compared to tobacco due to the variety of strain options and lack of controlled, research-grade cigarette products available. As such, we opted to establish our model using indica-dominant strains with THC levels of 10 – 14% and CBD levels of 0 – 2 % due to the popularity of this composition among recreational users^22^. At the time of this report, no licensed cannabis retailers in Canada manufacture identical cannabis cigarettes on a scale large enough to facilitate research. Thus, we developed an in-house method to produce consistent cigarettes using ground dried cannabis (Fig. 1). This in-house method is scalable to other research labs. In addition, we assessed both male and female mice, given the growing body of evidence that sex differences play an integral role in cannabis metabolism and cannabinoid signaling^33–35^. With these experimental variables implemented, our cannabis smoke exposure system effectively models real-world cannabis use patterns while respecting the need for reliability in a research environment.

Our cannabis smoke exposure model was effective at delivering cannabis smoke to mice. Within the exposure chamber, mice displayed common symptoms of smoke exposure seen in our previous tobacco studies but tolerated smoke exposure well^25,36^. While studies have demonstrated that cannabis smoke increases locomotion in rats^37^, our model appears to have the opposite effect, potentially due to the opposing effects of prolonged time in the smoke-filled exposure chamber. TPM in the chamber was seen to be at similar levels to those seen in tobacco smoke studies using the same system^25,36^, which may also contribute to the tobacco smoke-like behavioural phenotype we observed. In our post-exposure analysis, we found elevated levels of COHb, THC, and carboxy-THC across the sexes that decreased 60 minutes post exposure (Fig. 2, Supp. Fig. 2). These increases, along with the lack of CBD present, are similar to pharmacokinetic trends in human cannabis smokers post cigarette consumption^38^. With these findings, we developed a model system which resulted in the systemic delivery of smoke and cannabinoids at a magnitude equivalent with human smokers.

Maintaining immune homeostasis in the lungs is essential in supporting positive respiratory health outcomes. Previous studies have demonstrated a strong link between tobacco smoke exposure, chronic pulmonary inflammation, and the development of chronic obstructive pulmonary disease (COPD)^39^ – a relationship not yet fully explored in terms of cannabis smoke. Instigating this relationship is a smoke-induced modulation of immune cell populations and mediators leading to a chronic state of lung inflammation and tissue damage. In particular, innate immune cell populations such as lung macrophages and neutrophils have been found to be the main effector cells in the development of smoke-associated lung inflammation^25–27^. Given this paradigm, we initiated our characterization of cannabis smoke-induced alterations to respiratory immunity by investigating whether cannabis smoke exposure alters immune cell populations. Our findings show that 4 days of cannabis smoke exposure modulated innate immune cell populations in the airways and lung tissue (Fig. 3, 4, 5). Specifically, we found that macrophages were increased in the airways of female smoke exposed mice and the lung tissue of male smoke exposed mice. These increases were matched by decreases in monocyte populations, suggesting that monocyte activation and differentiation may be induced as a consequence of cannabis smoke exposure. In addition, our analysis of lung macrophage subpopulations demonstrates a potential effect on subtypes that have been associated with heightened inflammatory profiles and may contribute to the development of lung pathologies^28,30^. Conversely, our data shows that four days of cannabis smoke exposure did not lead to any changes in neutrophils in the airways or lung tissue. While the current tobacco smoke literature suggests that exacerbated neutrophilia in the lungs is a prominent contributor to smoke-associated chronic inflammation, increases in macrophage quantity and phenotypical changes are thought to be the driver of exacerbated neutrophilic infiltration into smoke-exposed lung tissue^26^. Therefore, our findings are consistent with the hypothesis that acute cannabis smoke exposure may be inducing the early population level immunomodulation that characterizes tobacco smoke-associated chronic inflammation.

**Figure 4.**
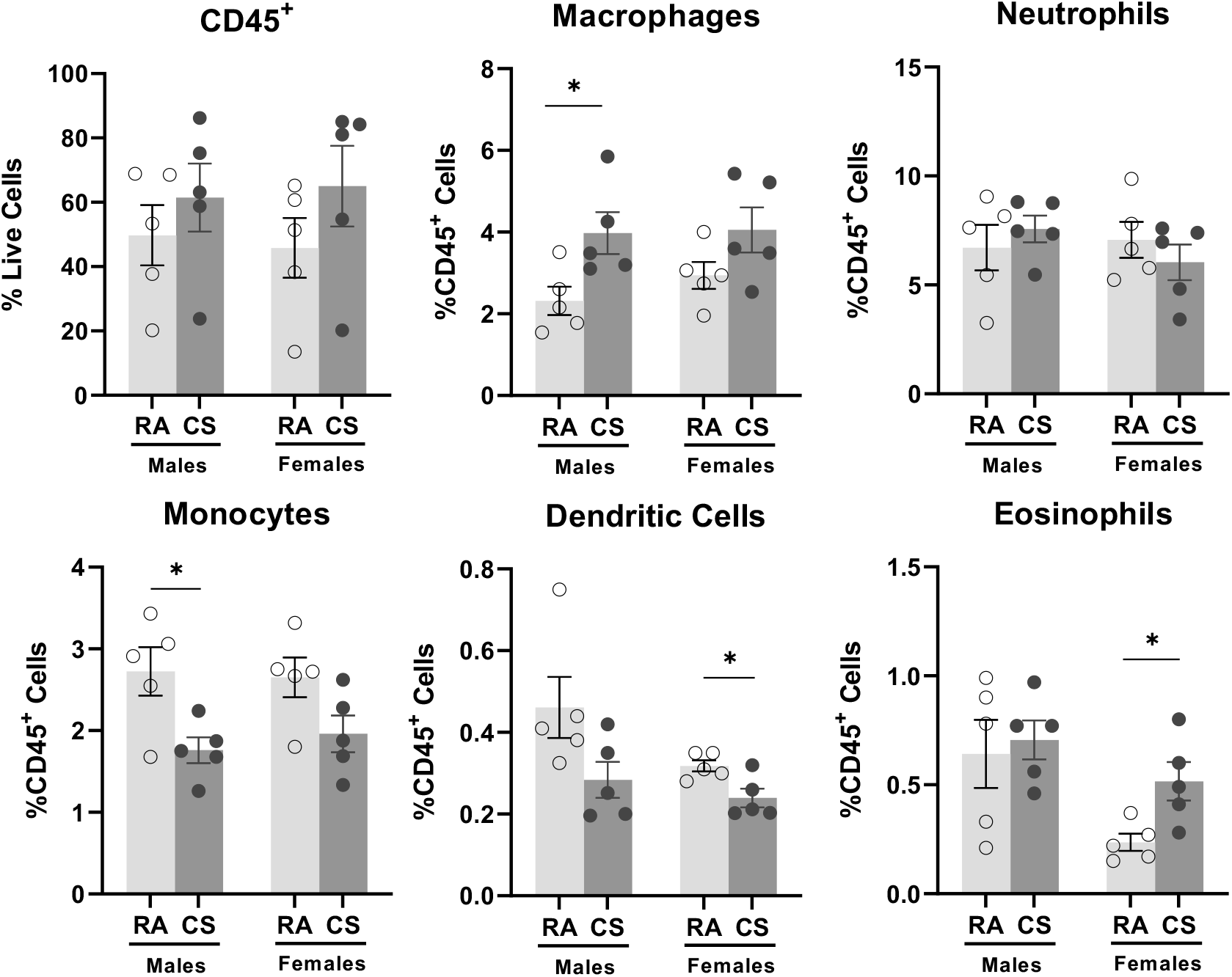
Cannabis smoke exposure modulates the proportionality of innate immune cell populations in the lungs. Male and female 6 – 8 week old BALB/c mice were exposed to room air (RA) or the smoke of 6 cannabis cigarettes (CS) twice a day for 4 days. Lung innate immune cell populations were quantified via flow cytometry. Proportionality was determined via haemocytometer. Data represent mean ± SEM; n = 5/group; *P<0.05, unpaired t-test within each sex.

Along with immune cell dysregulation, elevated proinflammatory cytokine levels have been associated with the induction of chronic lung inflammation and the development of COPD^40^. Tobacco and cannabis smoke has been shown to modulate immune mediators involved in anti-viral signaling^31,32^. Our immune mediator analysis demonstrates a subtle modulation of the detected cytokines and chemokines in the lung tissue after four days of cannabis smoke exposure (Fig. 5). Of the detected mediators, our findings indicated increases of macrophage-derived chemokine (MDC), CCL17 (TARC), and vascular endothelial growth factor (VEGF) in both sexes. MDC and TARC are both macrophage-associated mediators which have been previously demonstrated to be increased by tobacco smoke exposure^42^. Similarly, VEGF, which is highly expressed in the lung epithelium, has been shown to be greatly elevated by tobacco smoking^43^. Combined, these changes demonstrate further correlations between cannabis and tobacco smoke-induced effects on lung immunity. Conversely, mediators previously implicated in tobacco smoke investigations, such as IL-1α, were not altered. This suggests that cannabis smoke may have differential immunomodulator affects on the lung environment. As well, other macrophage and monocyte associated mediators involved in activation and differentiation were undetected or unchanged. Anti-viral signaling molecules such as IFNβ-1, IP-10, and RANTES were also unchanged. Consequently, the overall change to the inflammatory environment as a result of four days of cannabis smoke exposure was minimal but suggestive that longer exposure protocols are warranted to understand chronic cannabis use.

**Figure 5.**
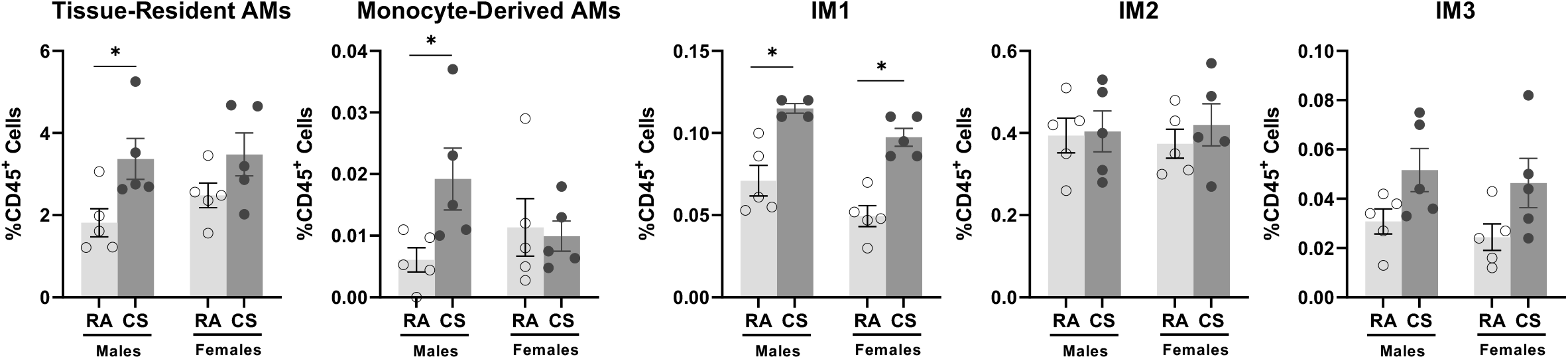
Cannabis smoke exposure modulates the proportionality of macrophage subpopulations in the lungs. Male and female 6 – 8 week old BALB/c mice were exposed to room air (RA) or the smoke of 6 cannabis cigarettes (CS) twice a day for 4 days. Macrophage subpopulations were quantified via flow cytometry. Proportionality was determined via haemocytometer. AM – Alveolar Macrophage, IM – Interstitial Macrophage. Data represent mean ± SEM; n = 4 – 5/group; *P<0.05, unpaired t-test within each sex.

**Figure 6.**
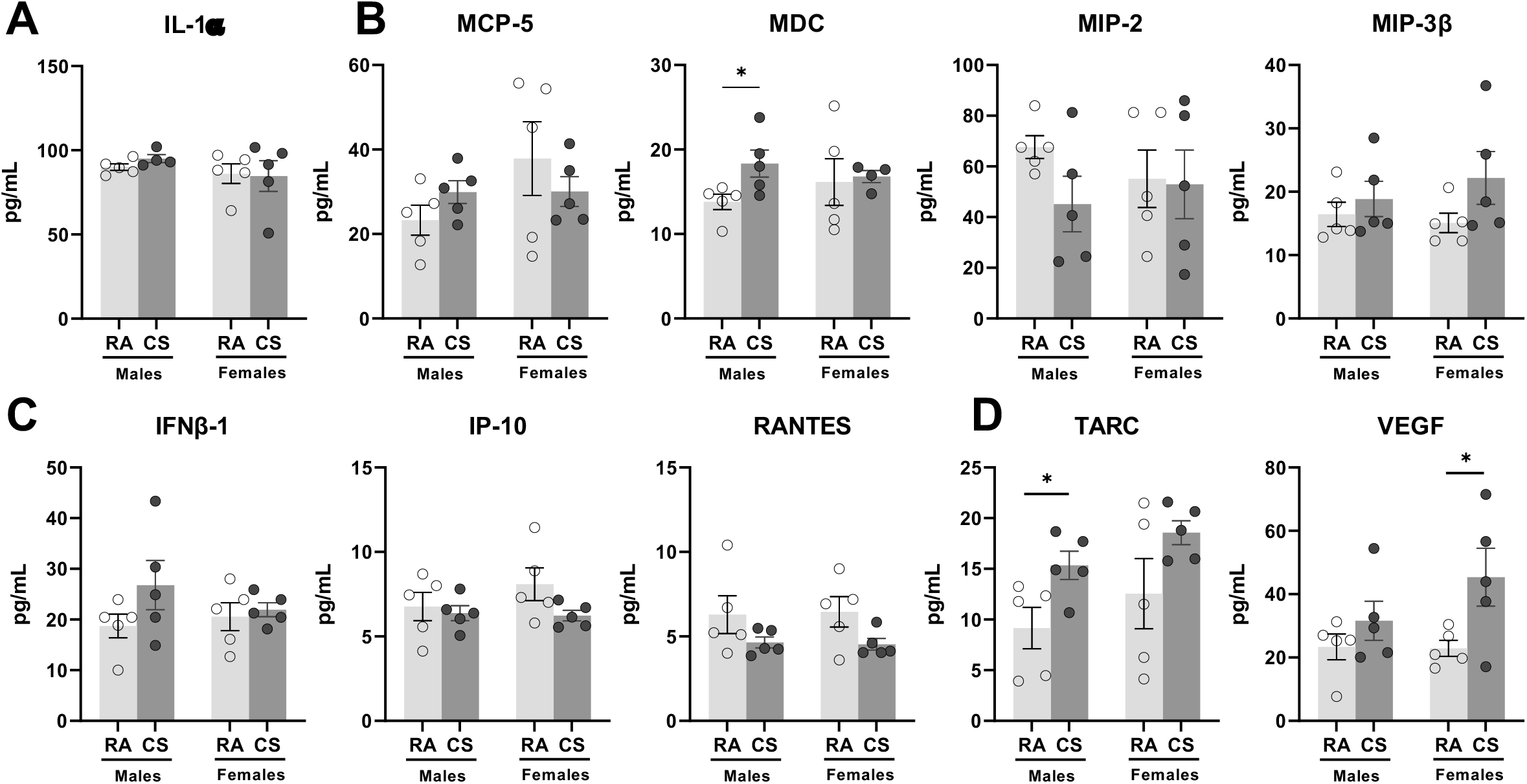
Impact of acute cannabis smoke exposure on immune mediators in the lungs. Male and female 6 – 8 week old BALB/c mice were exposed to room air (RA) or the smoke of 6 cannabis cigarettes (CS) twice a day for 4 days. Immune mediators were quantified via multiplex analysis (Eve Technologies). Shown mediators are those with quantities above the analysis’ lower level of detection and are associated with (A) smoke-associated inflammation, (B) macrophage cell signaling, and (C) anti-viral signaling. (D) Other immune mediators with significant differences above the lower level of detection are also shown. Data represent mean ± SEM; n = 5/group; *P<0.05, unpaired t-test within each sex.

Updated models of cannabis smoke exposure are needed to investigate how modern cannabis strains, which significantly vary from historical strains, impact lung health and disease. To address this unmet need, we characterized a model of smoke exposure, focusing on metabolic analysis for phytocannabinoids, and lung immune profiling, in both male and female mice. Our model system recapitulates the behavioural and pharmacokinetic changes observed in human cannabis smokers, providing evidence that the exposure protocol is within real-world ranges. Cannabis exposure resulted in innate immune cell changes that are proposed to precede more chronic inflammation in tobacco smokers using similar exposure systems. Our model lays a foundation for long-term exposures, which will add to our understanding of how immune cell populations and mediators are impacted by cannabis, and whether this contributes to adverse lung health outcomes. Additionally, in both acute and chronic scenarios, our model can be leveraged for smoke/pathogen co-exposures to interrogate the potential pathological consequences of the cannabis smoke-induced immune changes observed. Collectively, our results define a validated modern cannabis smoke exposure model essential for studying the relationship between cannabis consumption and respiratory health – an increasingly important undertaking given the growing popularity of cannabis on a global scale.

## Acknowledgements

The authors acknowledge Joanna Kasinska for her technical expertise and contributions to model development. Additionally, the authors acknowledge Dr. Tracey Campbell and Nikki Henriquez (McMaster University Centre for Microbial Chemical Biology) and Dr. Stephen Hill (McMaster University Medical Centre Core Laboratory) for their assistance in metabolite quantification. Lastly, Mr. Allen Fein is acknowledged for his oversight and execution of our cannabis research license through Health Canada.

---

**Supp. Figure 1.**
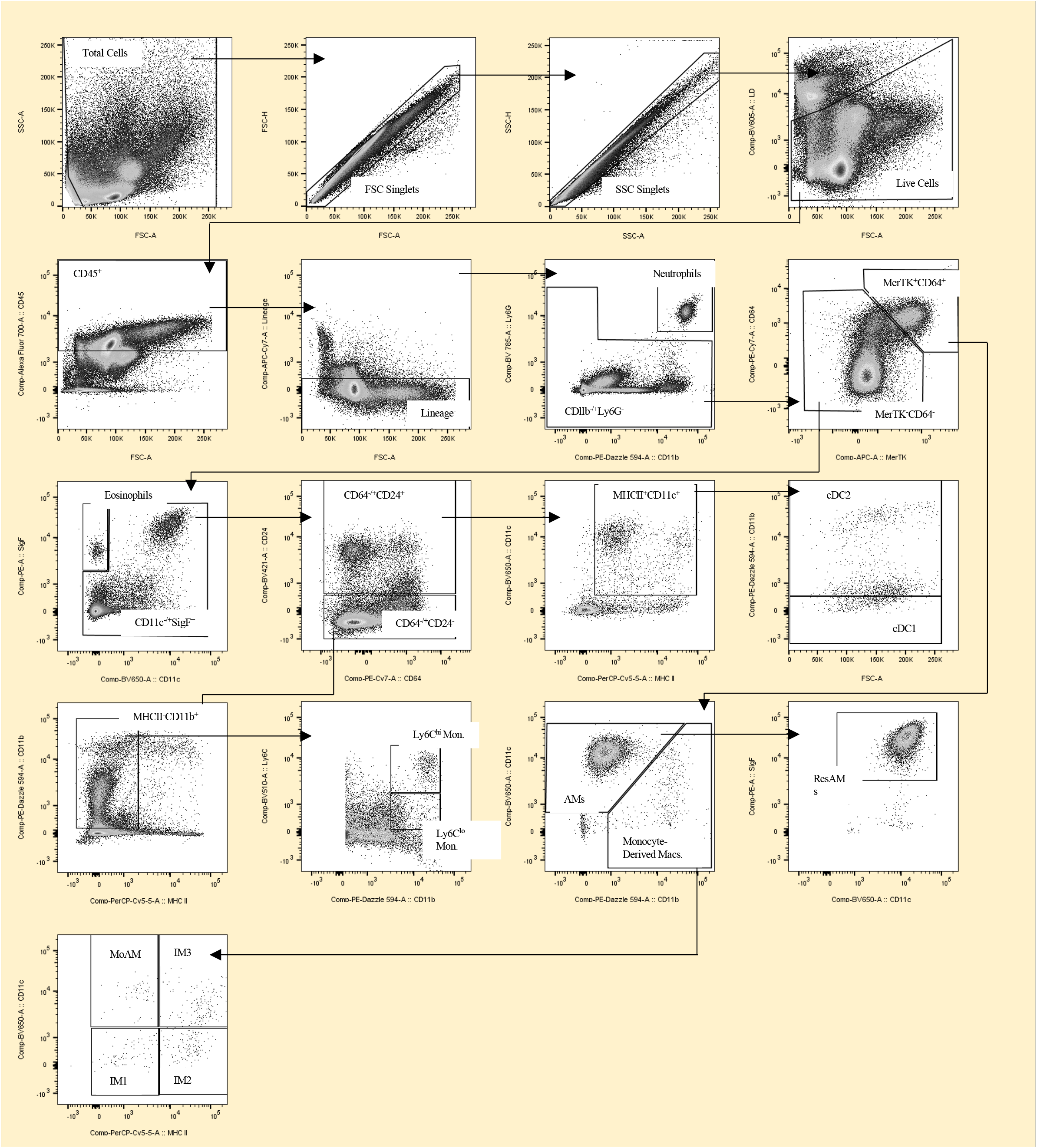
Lung tissue innate immune cell flow cytometry gating strategy. Gating was determined based on population identification and fluorescence-minus-one controls. Gating strategy generated using lung tissue collected from a room air exposed female control mouse. Lineage gate contains B220+, CD3e+, and EpCAM+ cell populations.

**Supp. Figure 2.**
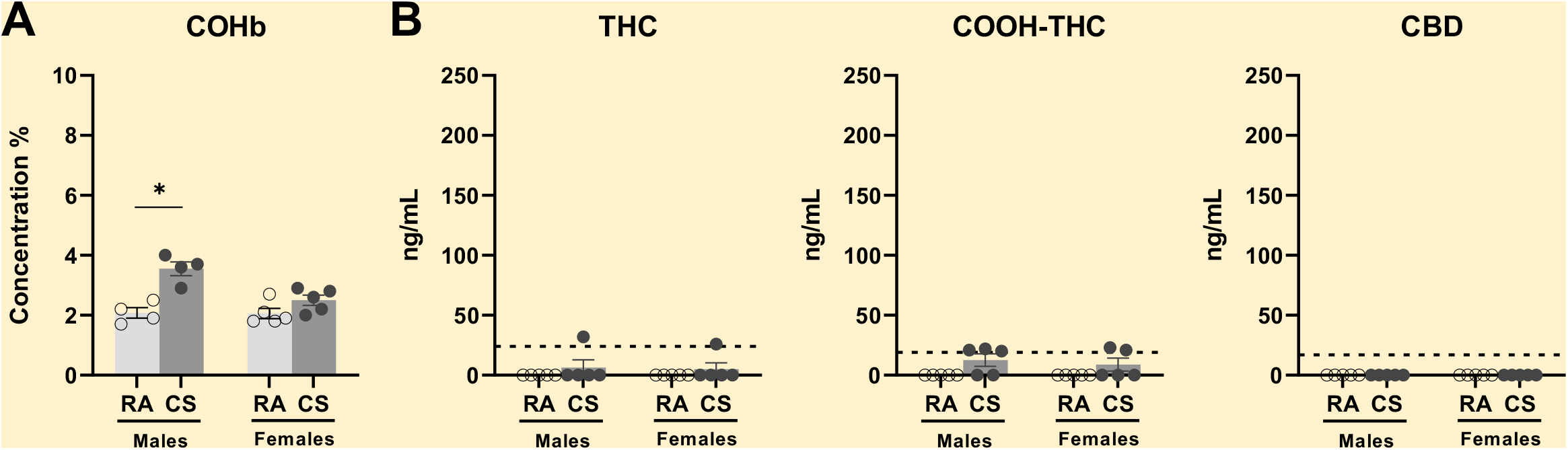
Acute cannabis smoke exposure increases plasma carboxyhemoglobin, THC, and carboxy-THC. Male and female 6 – 8 week old BALB/c mice were exposed to room air (RA) or the smoke of 6 cannabis cigarettes (CS) twice in a day. Whole blood was collected via retro-orbital bleed 60 minutes following the second exposure session. (A) Carboxyhemoglobin (COHb) percentage was quantified via CO-oximetry. (B) Plasma tetrahydrocannabinol (THC), cannabidiol (CBD), and carboxy-tetrahydrocannabinol (COOH-THC) were quantified via mass spectroscopy. Dotted lines represent the limit of detection for specific cannabinoid. Data points at 0 ng/mL represent values below the limit of detection in cannabinoid analysis. Data represent mean ± SEM; n = 3 – 5/group; *P<0.05, unpaired t-test within each sex in COHb analysis.

**Supp. Table 1.** Multiplex analysis of immune mediator expression in the lungs of male and female cannabis smoke exposed mice. Experimental and lower limit of detection (LLD) values are represented in pg/mL. Bolded factors have quantities above the LLD and contain statistical significance in at least one sex. n = 5/group.

|  |  | Males |  |  |  | Females |  |  |  |
| --- | --- | --- | --- | --- | --- | --- | --- | --- | --- |
| Factor | LLD | RA<br>(SEM) | CS<br>(SEM) | CS:RA<br>Ratio | P value | RA<br>(SEM) | CS<br>(SEM) | CS:RA<br>Ratio | P value |
| 6Ckine/Exodus2 | 4.00 | 15711.00<br>(3801.00) | 18142.00<br>(1530.00) | 1.15 | 0.57 | 21084<br>(3706.00) | 14610<br>(430.00) | 0.69 | 0.12 |
| Eotaxin | 0.40 | 33.74<br>(4.98) | 42.18<br>(2.56) | 1.25 | 0.17 | 41.49<br>(5.80) | 42.11<br>(2.58) | 1.01 | 0.93 |
| EPO | 8.00 | 0.87<br>(0.05) | 0.94<br>(0.08) | 1.08 | 0.61 | 0.83<br>(0.03) | 0.84<br>(0.06) | 1.01 | 0.87 |
| Fractalkine | 8.00 | 75.68<br>(15.25) | 80.21<br>(10.72) | 1.06 | 0.81 | 64.19<br>(13.62) | 69.15<br>(9.74) | 1.08 | 0.77 |
| G-CSF | 0.50 | 1.00<br>(0.18) | 0.79<br>(0.08) | 0.79 | 0.37 | 1.06<br>(0.13) | 1.54<br>(0.50) | 1.45 | 0.38 |
| GM-CSF | 10.90 | 2.26<br>(0.17) | 2.42<br>(0.10) | 1.07 | 0.43 | 2.23<br>(0.06) | 2.05<br>(0.09) | 0.92 | 0.12 |
| IFNβ-1 | 10.00 | 18.74<br>(2.33) | 26.79<br>(4.86) | 1.43 | 0.17 | 20.58<br>(2.75) | 21.93<br>(1.36) | 1.07 | 0.67 |
| IFNγ | 0.40 | 4.37<br>(0.19) | 4.87<br>(0.30) | 1.11 | 0.15 | 3.63<br>(0.80) | 4.19<br>(0.60) | 1.15 | 0.59 |
| IL-1α | 10.30 | 90.03<br>(1.94) | 95.09<br>(2.40) | 1.06 | 0.14 | 86.10<br>(5.79) | 84.71<br>(9.15) | 0.98 | 0.90 |
| IL-1β | 5.40 | 1.25<br>(0.18) | 2.94<br>(0.20) | 2.35 | <0.01 | 1.67<br>(0.51) | 1.37<br>(0.48) | 0.82 | 0.68 |
| IL-2 | 0.60 | 6.06<br>(0.81) | 10.32<br>(2.16) | 1.70 | 0.10 | 6.28<br>(0.60) | 6.43<br>(1.20) | 1.02 | 0.91 |
| IL-3 | 0.70 | 0.51<br>(0.01) | 0.48<br>(0.01) | 0.94 | 0.17 | 0.49<br>(0.02) | 0.50<br>(0.02) | 1.02 | 0.79 |
| IL-4 | 0.20 | 0.46<br>(0.01) | 0.48<br>(0.01) | 1.04 | 0.14 | 0.47<br>(0.01) | 0.47<br>(0.01) | 1.00 | >0.99 |
| IL-5 | 0.30 | 0.45<br>(0.01) | 0.50<br>(0.06) | 1.11 | 0.40 | 0.46<br>(0.01) | 0.43<br>(0.01) | 0.93 | 0.14 |
| IL-6 | 0.40 | 0.47<br>(0.06) | 0.58<br>(0.09) | 1.23 | 0.34 | 0.50<br>(0.05) | 0.41<br>(0.01) | 0.82 | 0.16 |
| IL-7 | 1.40 | 1.36<br>(0.20) | 1.97<br>(0.56) | 1.45 | 0.32 | 1.51<br>(0.16) | 1.68<br>(0.30) | 1.11 | 0.64 |
| IL-9 | 17.30 | 7.50<br>(1.90) | 9.74<br>(3.16) | 1.30 | 0.56 | 9.05<br>(1.80) | 8.53<br>(1.37) | 0.94 | 0.82 |
| IL-10 | 1.20 | 0.43<br>(0.04) | 0.47<br>(0.08) | 1.09 | 0.62 | 0.66<br>(0.12) | 0.47<br>(0.16) | 0.71 | 0.36 |
| IL-11 | 10.00 | 2.24<br>(0.24) | 2.39<br>(0.31) | 1.07 | 0.71 | 2.74<br>(0.38) | 2.39<br>(0.18) | 0.87 | 0.42 |
| IL-12p40 | 3.90 | 1.61<br>(0.95) | 3.52<br>(2.20) | 2.19 | 0.45 | 0.61<br>(0.33) | 0.08<br>(0.06) | 0.13 | 0.17 |
| IL-12p70 | 4.80 | 2.10<br>(0.11) | 1.78<br>(0.10) | 0.85 | 0.06 | 1.89<br>(0.06) | 1.79<br>(0.08) | 0.95 | 0.38 |
| IL-13 | 33.30 | 1.28<br>(0.03) | 1.31<br>(0.02) | 1.02 | 0.35 | 1.30<br>(0.02) | 1.30<br>(0.04) | 1.00 | 0.88 |
| IL-15 | 7.40 | 5.41<br>(0.59) | 4.78<br>(0.69) | 0.88 | 0.51 | 3.88<br>(0.37) | 4.75<br>(0.49) | 1.22 | 0.20 |

**Supp. Table 1. Multiplex analysis of immune mediator expression in the lungs of male and female cannabis smoke exposed mice.**
|  |  | Males |  |  |  | Females |  |  |  |
| --- | --- | --- | --- | --- | --- | --- | --- | --- | --- |
| Factor | LLD | RA<br>(SEM) | CS<br>(SEM) | CS:RA<br>Ratio | P value | RA<br>(SEM) | CS<br>(SEM) | CS:RA<br>Ratio | P value |
| IL-16 | 1.00 | 3232.00<br>(184.00) | 3034.00<br>(243.80) | 0.94 | 0.54 | 3255.00<br>(458.70) | 3148.00<br>(316.00) | 0.97 | 0.85 |
| IL-17 | 0.20 | 0.51<br>(0.01) | 0.54<br>(0.03) | 1.06 | 0.29 | 0.54<br>(0.02) | 0.52<br>(0.02) | 0.96 | 0.41 |
| IL-20 | 21.00 | 2.75<br>(0.46) | 2.54<br>(0.17) | 0.92 | 0.69 | 3.07<br>(0.44) | 4.18<br>(0.19) | 1.36 | 0.07 |
| IP-10 | 0.10 | 6.77<br>(0.84) | 6.38<br>(0.44) | 0.94 | 0.69 | 8.09<br>(0.97) | 6.24<br>(0.31) | 0.77 | 0.11 |
| KC | 1.80 | 12.20<br>(1.63) | 13.99<br>(1.67) | 1.15 | 0.46 | 10.71<br>(2.53) | 7.27<br>(2.55) | 0.68 | 0.37 |
| LIF | 0.50 | 0.55<br>(0.02) | 0.48<br>(0.03) | 0.87 | 0.11 | 0.54<br>(0.03) | 0.47<br>(0.01) | 0.87 | 0.06 |
| LIX | 22.10 | 3.43<br>(0.12) | 3.57<br>(0.38) | 1.04 | 0.74 | 3.53<br>(0.31) | 3.73<br>(0.39) | 1.07 | 0.70 |
| MCP-1 | 6.70 | 1.92<br>(0.13) | 1.98<br>(0.37) | 1.03 | 0.89 | 2.43<br>(0.10) | 1.83<br>(0.08) | 0.75 | <0.01 |
| MCP-5 | 1.00 | 23.29<br>(3.54) | 29.92<br>(2.71) | 1.28 | 0.17 | 37.85<br>(8.74) | 30.06<br>(3.53) | 0.79 | 0.43 |
| M-CSF | 3.50 | 0.19<br>(0.01) | 0.21<br>(0.04) | 1.11 | 0.71 | 0.38<br>(0.06) | 0.14<br>(0.02) | 0.37 | 0.02 |
| **MDC** | **2.00** | **13.80**<br>**(0.91)** | **18.34**<br>**(1.61)** | **1.33** | **0.04** | **16.15**<br>**(2.76)** | **16.81**<br>**(0.71)** | **1.04** | **0.84** |
| MIG | 2.40 | 18.33<br>(2.97) | 23.17<br>(4.27) | 1.26 | 0.38 | 32.97<br>(8.60) | 31.00<br>(4.00) | 0.94 | 0.84 |
| MIP-1α | 7.70 | 2.00<br>(0.35) | 3.97<br>(1.22) | 1.99 | 0.16 | 2.15<br>(0.30) | 1.76<br>(0.28) | 0.82 | 0.38 |
| MIP-1β | 11.90 | 0.38<br>(0.05) | 0.28<br>(0.06) | 0.74 | 0.24 | 0.31<br>(0.05) | 0.43<br>(0.13) | 1.39 | 0.39 |
| MIP-2 | 30.60 | 67.61<br>(4.53) | 45.16<br>(10.98) | 0.67 | 0.10 | 55.15<br>(11.34) | 52.94<br>(13.53) | 0.96 | 0.90 |
| MIP-3α | 5.00 | 1.05<br>(0.07) | 1.35<br>(0.25) | 1.29 | 0.29 | 1.09<br>(0.12) | 1.09<br>(0.10) | 1.00 | 0.97 |
| MIP-3β | 8.00 | 16.44<br>(1.91) | 18.87<br>(2.79) | 1.15 | 0.49 | 15.07<br>(1.53) | 22.17<br>(4.17) | 1.47 | 0.15 |
| RANTES | 2.70 | 6.29<br>(1.12) | 4.65<br>(0.33) | 0.74 | 0.20 | 6.46<br>(0.90) | 4.53<br>(0.34) | 0.70 | 0.08 |
| **TARC** | **2.00** | **9.15**<br>**(2.04)** | **15.34**<br>**(1.40)** | **1.68** | **0.04** | **12.56**<br>**(3.45)** | **18.56**<br>**(1.18)** | **1.48** | **0.14** |
| TIMP-1 | 2.00 | 223.70<br>(23.69) | 251.60<br>(20.44) | 1.12 | 0.40 | 190.60<br>(43.24) | 207.50<br>(9.30) | 1.09 | 0.71 |
| TNFα | 1.70 | 0.11<br>(0.01) | 0.11<br>(0.01) | 1.00 | 0.60 | 0.11<br>(0.01) | 0.11<br>(0.01) | 1.00 | 0.85 |
| **VEGF** | **0.10** | **23.36**<br>**(4.07)** | **31.63**<br>**(6.17)** | **1.35** | **0.30** | **22.90**<br>**(2.50)** | **45.39**<br>**(9.13)** | **1.98** | **0.04** |

## Notes

Funding: This work was supported by the Ontario Lung Association and the Michael G. DeGroote Centre for Medicinal Cannabis Research

### Competing Interest Statement

The authors have declared no competing interest.

